# A mathematical model of tumor growth driven by oxygen–lactate–exosome feedback

**DOI:** 10.64898/2026.10.10.758176

**Authors:** Gopinath Sadhu, D C Dalal, Helen M. Byrne

**Affiliations:** Department of Mathematics, Indian Institute of Technology Guwahati, Guwahati, India; Wolfson Centre for Mathematical Biology, Mathematical Institute, University of Oxford, Oxford, United Kingdom; Ludwig Institute for Cancer Research, University of Oxford, Oxford, United Kingdom; Department of Bioengineering, Indian Institute of Science, Bangalore, Karnataka, India

**Author notes:** Corresponding author *Email address:* (Gopinath Sadhu).

**Keywords:** Lactate, exosome, tumor growth, mathematical model

## Abstract

Solid tumors develop within a heterogeneous microenvironment shaped by coupled biochemical and metabolic processes. As a tumor grows, oxygen gradients develop from the periphery towards the core, giving rise to hypoxic regions. Hypoxia induces a metabolic shift from oxidative phosphorylation to glycolysis, leading to elevated lactate production. Experimental studies further indicate that increased lactate levels enhance the secretion of tumor-derived exosomes carrying proteins, mRNA, and lipids, which in turn promote tumor cell proliferation. Together, these interactions form a feedback loop linking oxygen availability, lactate accumulation, and exosome-mediated signaling. Despite growing experimental evidence for each component, the impact of this coupled oxygen–lactate–exosome axis on tumor growth dynamics has not yet been systematically explored from a mathematical perspective. It remains unclear how the oxygen-lactate-exosome axis affects tumor growth - impact of nonlocal effects.

In this paper, we develop a mechanistic mathematical model to investigate how this feedback mechanisms affects tumor growth. We first consider an in vitro setting oxygen levels are spatially homogeneous and vary periodically in time between normoxic and hypoxic states, capturing key features of cyclic hypoxia observed in vivo. Numerical simulations reveal that exosome-mediated feedback has a pronounced effect on tumor volume under chronic hypoxia. In addition, we observe that the tumor volume decreases as the period of cyclic hypoxia increases. We then extend the model to an in vivo framework by incorporating oxygen diffusion and consumption, allowing spatial heterogeneity to emerge. Simulations reveal that tumor cells in normoxic regions benefit indirectly from enhanced exosome secretion driven by lactate produced in hypoxic regions. Overall, the results show how spatially mediated metabolic signaling can amplify tumor growth and aggressiveness, and illustrate how mathematical modeling can increase understanding of non-local feedback mechanisms within the tumor microenvironment.

## 1. Introduction

Tumor growth is a complex biological process that begins when a genetically altered cell acquires the ability for uncontrolled, indefinite proliferation [1]. This leads to the formation of a cell cluster that expands over time, receiving essential nutrients and oxygen from pre-existing blood vessels. Such a mass is referred to as an avascular tumor. The limited diffusion distance from blood vessels to the tumor site, combined with high oxygen consumption at the tumor periphery, creates an oxygen gradient in the tumor central region. Consequently, a hypoxic core emerges when oxygen levels fall below a critical threshold, prompting a metabolic shift from oxidative phosphorylation to glycolysis. This shift leads to excessive lactate production, which acidifies the tumor microenvironment [2]. Elevated lactate concentrations are toxic to surrounding healthy cells, contribute to extracellular matrix (ECM) degradation, and function as immune suppressors [3, 4]. Lactate also upregulates the secretion of tumor-derived exosomes, which serve as carriers of lipids, proteins, and mRNAs [5]. Using exosomes, tumor cells communicate with neighboring cells and also distant parts of the primary tumor. The roles of exosomes are not limited to serving as messengers; they also participate in various oncogenic processes, including immune evasion, angiogenesis, collective cell migration, and epithelial-to-mesenchymal transition (EMT) [6, 7, 8]. Experimental studies reported that exosomes increase tumor cell proliferation [9]. Logozzi et al. [5] recently demonstrated that tumor-derived exosome secretion increases in the presence of lactate. The relationship between lactate and hypoxia is well established in tumor biology. However, the mechanisms by which tumor growth progresses along the oxygen-lactate-exosome axis remain poorly understood.

Numerous mathematical models have been developed to elucidate tumor growth dynamics [10, 11, 12]. These models are generally classified into three categories: continuum, discrete, and hybrid frameworks. In continuum models, biological entities such as cells and chemical concentrations are represented as continuous variables. In contrast, discrete models treat entities as individual units, which enables a more accurate representation of cellular interactions. In a hybrid modeling framework, the multiscale nature of biological phenomena is accurately captured through a mix of continuum and discrete techniques. Many researchers used ordinary differential equations (ODEs) to model population-level interactions among different cell types, such as heterogeneous tumor cells, vascular cells, immune cells, and dead cells [13, 14, 15]. In addition, the temporal dynamics of tumor growth under nutrient conditions and the effects of radiation therapy are well explored [16, 17]. Tumor growth processes are not readily captured by an ODE-based model, as they are intertwined with spatial interactions within the tumor’s surrounding medium, nutrient gradients, and oxygen heterogeneity. Several spatio-temporal models have been proposed using partial differential equations (PDEs) to elucidate the spatial effects. In 1972, a seminal work by Greenspan [18] presented a mathematical model of tumor growth limited by nutrient diffusion. This approach has been adopted by many other researchers to study the effects of hypoxia and the microenvironment on tumor progression [19, 20]. Under hypoxic stress, tumor cells switch from oxidative phosphorylation to glycolysis for glucose metabolism. As a result, lactate, H+ ions, and CO_2_ are produced in ample amounts, which acidify the tumor microenvironment. Casciari et al. [20] used a mathematical model to investigate the effects of tumor microenvironmental acidity on its growth dynamics. Lactate is not a metabolic waste; it contributes to energy production in the normoxic region, as it is consumed by normoxic cells [21]. Sadhu and Dalal [22] proposed a mathematical model to elucidate how oxygen and lactate interplays in a tumor microenvironment under the above biological phenomenon. Also, they studied effects of tumor growth dynamics on hypoxia-lactate axis [23]. Tumor cells secrete various growth factors to accelerate their progression. Additionally, tumor-derived exosomes promote cellular proliferation. Lai and Friedman [24] developed a mathematical model to elucidate the influence of exosomal miRNA on lung cancer progression. They subsequently applied a similar modeling framework to investigate the effects of microRNA in colorectal cancer [25]. Exosome secretion is influenced by the lactate status of the tumor microenvironment, which is closely linked to local oxygen concentration. The interactions among tumor cells, oxygen, lactate, and exosomes remain poorly understood.

In this article, we present a continuum mathematical model to investigate changes in tumor growth dynamics along the oxygen-lactate-exosome axes. The model comprises a coupled system of partial differential equations (PDEs) describing the concentrations of oxygen, lactate, and exosomes, as well as the densities of tumor and necrotic cells. The model assumes that lactate production and consumption rates depend on local oxygen levels, and that exosome secretion rates increase in the presence of lactate. Additionally, tumor cell proliferation is assumed to increase with higher exosome and oxygen concentrations, while the tumor cell death rate rises with lactate accumulation and decreasing oxygen levels. The numerical results begin with two control scenarios: oxygen-rich and oxygen-deprived conditions, in which oxygen concentration is distributed homogeneously throughout the tumor domain. Oxygen levels within the tumor site are not constant in space or time. Recent experimental evidence indicates that oxygen levels may fluctuate between oxygen-rich and oxygen-deprived states across a wide range of timescales, from minutes to days, a phenomenon known as cyclic hypoxia [23, 26]. The current study also examines the impact of cyclic hypoxia on tumor growth dynamics within the oxygen-lactate-exosome framework. Additionally, oxygen levels in the tumor exhibit spatial heterogeneity due to rapid consumption by tumor cells and diffusion from primary blood vessels into the solid tumor region [27]. Moreover, we investigate how spatial oxygen heterogeneity influences tumor growth progression along the oxygen-lactate-exosome axis.

### 1.1. Article layout

The remainder of this paper is organized as follows. In Section 2 we introduce the proposed mathematical model and in Section 3 we nondimensionalize it, introduce typical parameter values and outline the numerical method used to generate model solutions (for details, see Appendix A). Simulation results are presented in Section 4. The paper concludes in 5 where we discuss our findings and outline possible directions for future work.

## 2. Mathematical model

In this section, we present our mathematical model for tumor growth driven by oxygen-lactate-exosome feedback based on the schematic diagram, given in Figure 1. It shows that tumor cells secrete exosomes, and this secretion is unregulated by the lactate produced by the tumor cells. Also, tumor cells’ proliferation is unregulated by tumor-derived exosomes.

**Figure 1:**
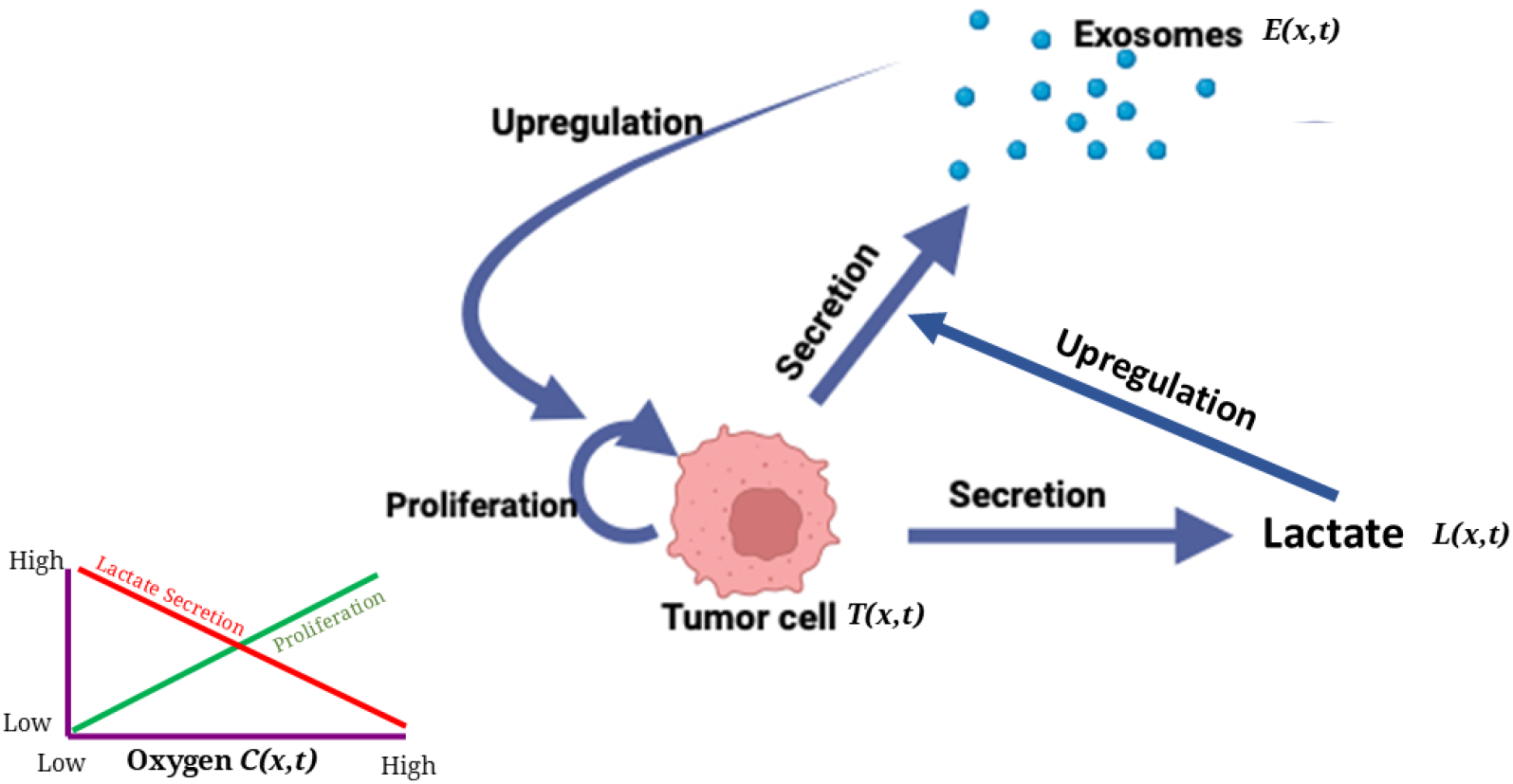
Schematic illustration of the interactions between tumor cells, exosomes, lactate, and oxygen availability in the tumor microenvironment. Tumor cells *T* (*x, t*) proliferate and secrete exosomes *E*(*x, t*) and lactate *L*(*x, t*). Exosomes further promote tumor progression through positive feedback upregulation. The inset depicts the dependence of tumor cell proliferation and lactate secretion on oxygen concentration *C*(*x, t*): proliferation increases with oxygen availability, whereas lactate secretion is enhanced under hypoxic (low-oxygen) conditions. This picture is created in Biorender.

**Figure 2:**
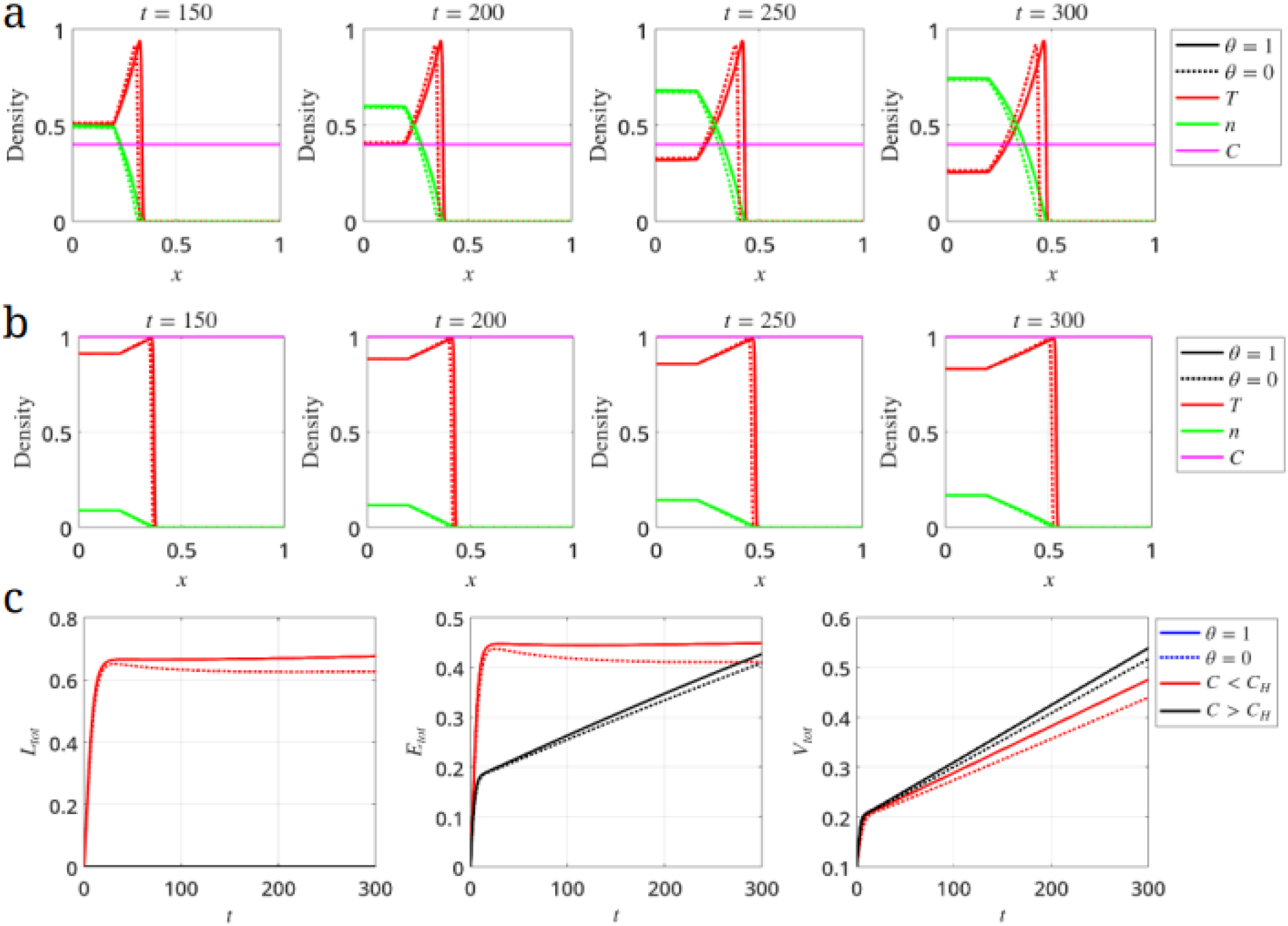
The spatial distribution of tumor cell density, necrotic cell density for oxygen deprived condition i.e., *C < C*_*H*_ in (a) and oxygen-rich condition i.e., *C > C*_*H*_ in (b). (c)Time evolution of total lactate, exosome, and tumor density in oxygen-rich and oxygen-deprived conditions.

For simplicity, we formulate our mathematical model in a one-dimensional Cartesian geometry. We use the principle of mass balance to derive partial differential equations (PDEs) that describe the spatio-temporal evolution of oxygen (*C*′(*x*′, *t*′)), lactate (*L*′(*x*′, *t*′)), exosomes (*E*′(*x*′, *t*′)), tumor cell density (*T* ′(*x*′, *t*′)) and dead cell density (*n*′(*x*′, *t*′)), where 0 *≤ x*′ *≤ R*′ denotes the spatial position and *t*′ *≥* 0 represents time.

### 2.1. Governing equations

#### Oxygen

We assume that the dominant processes regulate oxygen levels within the domain are: diffusion, consumption by tumor cells and natural decay. We assume that tumor cells consume oxygen at a rate that is an increasing, saturating function of *C*′. Combining these processes, we deduce that the following reaction-diffusion equation (RDE) governs the oxygen dynamics:

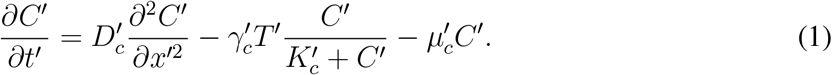

In Eq. (1), 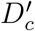 denotes the oxygen diffusion coefficient, 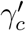 is the maximum rate at which tumor cells consume oxygen, 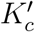 is the oxygen concentration at which consumption is half-maximal, and 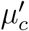 is the rate at which oxygen undergoes natural decay.

#### Lactate

We assume that the dominant processes governing lactate are: diffusion, production by tumor cells under hypoxia (where 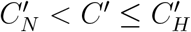), uptake by tumor cells under normoxia (where 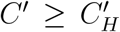), and natural decay. Combining these processes, we arrive at the following RDE for lactate:

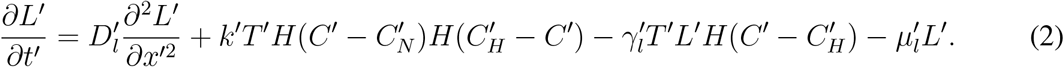

In Eq. (2), 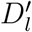 represents the diffusion coefficient of lactate, 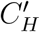 is the threshold oxygen concentration at which tumor cells become hypoxic and 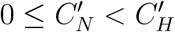 is the threshold oxygen concentration at which they undergo necrotic cell death. Finally, the non-negative parameter 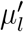 represents the rate at which lactate undergoes natural decay.

#### Exosomes

We consider that the dominant processes deriving exosomes level with the tumor domain are: diffusion, secretion by tumor cells and natural decay. Here, we assume that exosome secretion increases with an increase in lactate in the tumor microenvironment (TME). It is modeled using Michaelis–Menten kinetic formulation. Exosomes may degrade in many ways, such as phagocytosis and clearance through circulatory systems. Combining all these processes, we obtain the following RDE for exosomes:

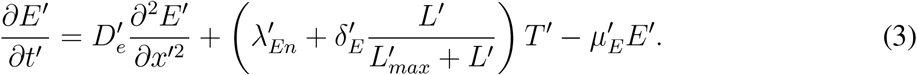

In Eq. (3), 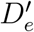 represents the diffusivity of exosomes, 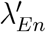 denotes the constant rate of exosome secretion by tumor cells, and 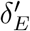 represents the rate of exosome secretion that is increasing, saturation function of *L*′. Here, 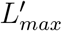 is the lactate concentration at which production is half-maximal, and 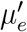 is the rate at which exosome undergoes natural decay.

#### Tumor cell

We model the spatio-temporal evolution of tumor cell density using the following RDE:

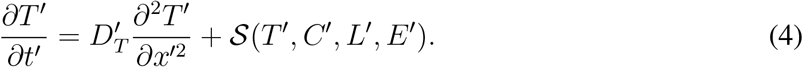

In Eq. (4), 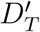 denotes the diffusion coefficient of tumor cells, and *S*(*T*′, *C*′, *L*′, *E*′) characterizes the tumor cells’ birth and death depend on oxygen (*C*′), lactate (*L*′) and exosomes (*E*′) concentrations. With an increased value of *C*′, cell proliferation increases. Cells start to die with an increased value of *L*′ after the critical value 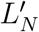. Also, exosomes contribute to the enhancement of tumor cell proliferation. Therefore, the functional form of *S*(*T*′, *C*′, *L*′, *E*′, *n*′) is given as

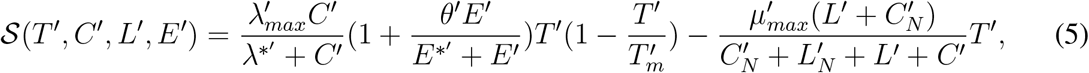

where the first term on RHS of Eq. (5) denotes the oxygen- and exosome-dependent logistic proliferation of tumor cells. Here, 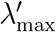 is the maximum proliferation rate of tumor cells, 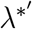 is the oxygen half-saturation constant, *θ*′ represents the maximum exosome-mediated enhancement of proliferation, and 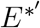 denotes the exosome half-saturation constant. The carrying capacity term 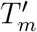 limits tumor growth due to space and resource constraints. The second term on the RHS of Eq. (5) denotes the lactate- and oxygen-dependent death rate of tumor cells. Here, 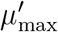 is the maximum death rate, while 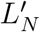 and 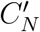 are the corresponding normalization constants regulating the relative contribution of lactate and oxygen to tumor cell death.

#### Necrotic core

We consider that the necrotic core forms through the lactate- and oxygen-dependent death of tumor cells and they undergo natural decay with a rate 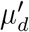. Combining these processes, we deduce the following dynamics equation for necrotic core:

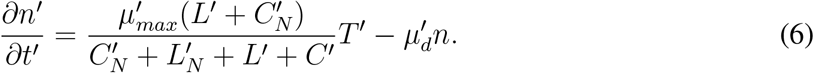

Taken together, Eqs. (1)-(6) describe how interactions between oxygen, lactate and exosome affects the tumor’s spatial composition and its growth dynamics.

### 2.2. Boundary and initial conditions

We close the model by imposing appropriate boundaries and initial conditions. We assume that the domain contains a tumor centered at *x*′ = 0 and that the system is symmetric about *x*′ = 0. Accordingly, we impose no-flux boundary conditions for oxygen concentration (*C*′), lactate concentration (*L*′), exosome concentration (*E*′), and tumor cell density (*T* ′) at *x*′ = 0:

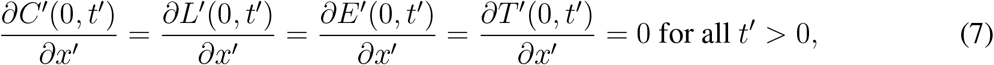

We assume further that lactate, exosome, and tumor cells do not leave the domain [0, *R*′], and that oxygen is maintained at constant value at the outer boundary. Thus, we have

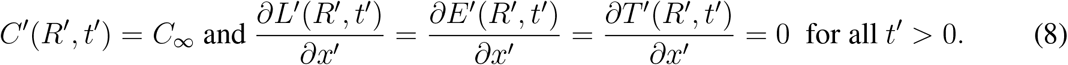

We assume that initially the oxygen distribution is spatially uniform across the domain, being maintained at a concentration consistent with the Eq. (8). In particular, we suppose that

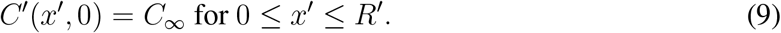

We suppose that the domain is initially devoid of lactate, so that

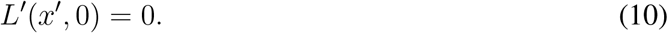

We assume that the tumor cells are initially uniformly distributed at concentration 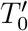 in a region and half-width *α*′ centered at *x*′ = 0. The initial distribution of exosome is assumed to have a similar functional form. Hence, these prescribe as

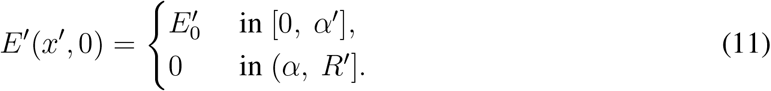

where 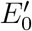 is the initial concentration of exosomes in the tumor tissue.

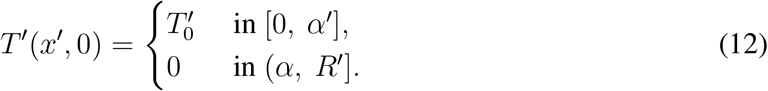

Finally, we suppose that the domain is initially devoid of necrotic tumor cells, so that

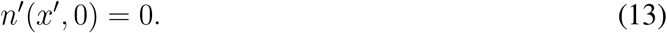

## 3. Non-dimensionalization

Before continuing, we initially non-dimensionalise the governing equations, scaling the independent and dependent variables with typical values in the following way:

- The reference length 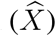 is the maximum length a tumor can invade 1 *cm* [28].
- The reference time scale, 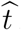 is the time scale for oxygen diffusion in the domain, i.e., 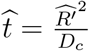, where *D*_*c*_ is the oxygen diffusion coefficient. Fixing *D*_*c*_ *≈* 2 *×* 10^*™*5^ *cm*^2^*s*^*™*1^ [29] and *R*′ = 1 *cm*, we estimate 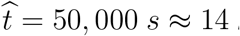 *hours*.
- Reference tumor cell density 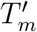, oxygen concentration *C*_*∞*_, lactate concentration 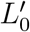 and exosome concentration 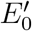 (where 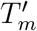 is maximum tumor cells capacity and its value 10^8^ *cells cm*^*™*3^, *C*_*∞*_ = 10 *mmHg* is the oxygen concentration is supplied at boundary. 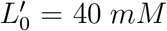 is the maximum lactate concentration in tumor medium. 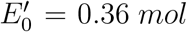 is the exosome concentration in physiological condition.

Under these scalings, the dimensionless variables are defined as,

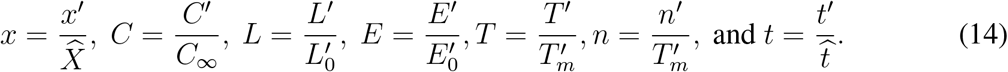

and the governing equations transform to give

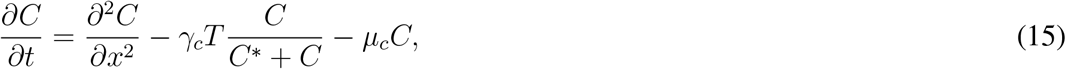

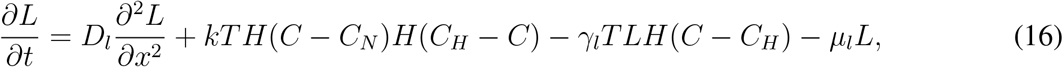

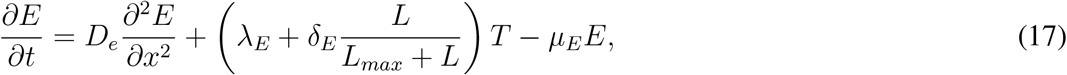

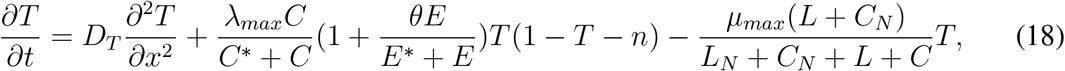

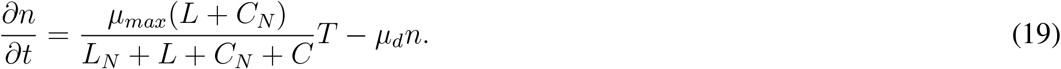

where the dimensionless parameter grouping are defined as 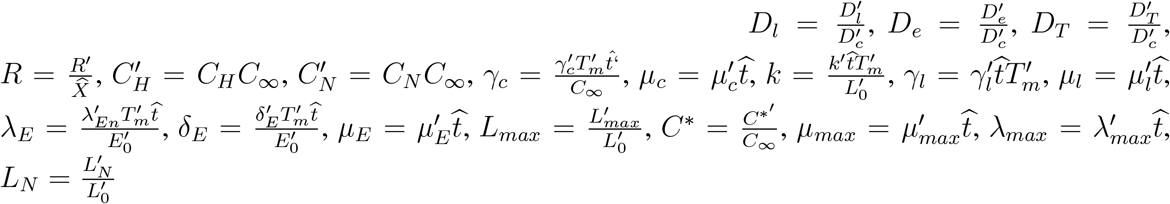 and 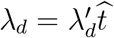.

### 3.1. Dimensionless boundary and initial conditions

The boundary and initial transform as follows:

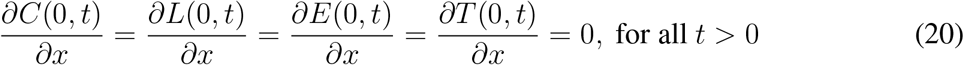

and

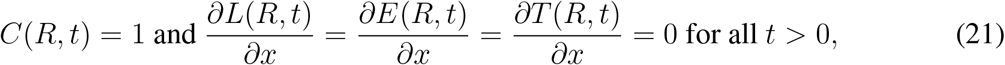

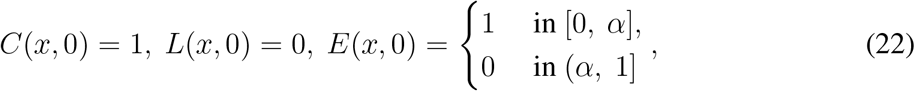

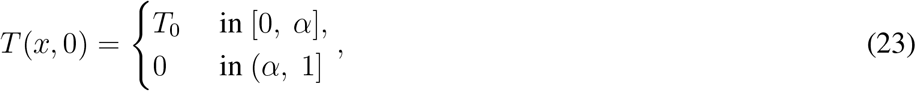

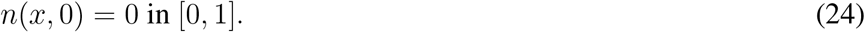

### 3.2. Parameter values

Most model parameters are estimated from the literature and are stated in Table 1. However, to the best of our knowledge, estimates and the parameters 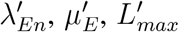 and 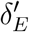 are not available in the literature. To estimate these parameters, we replicated the *in vitro* conditions used by Luo et al. [30]. They exposed melanoma (B16-F10) and liver (Hepa1-6) cells to fixed concentration of lactate with a constant lactate concentration to investigate its effects on exosome accumulation in the TME. After culturing 3 *×* 10^5^ cells for 24 hours in lactate concentrations of *L*′ = 0, 2, 5, and 10 *mM*, they measured exosome levels using flow cytometry. In the Table B.3, we summerise the normalized estimate that they obtained. We have added a detail description how 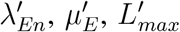 and 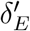 were obtained in Appendix B.

**Table 1:** Dimensional parameter values.

| Parameter | Value | Source |
| --- | --- | --- |
| $D_l'$ | $0.432 \text{ cm}^2 \text{ day}^{-1}$ | [3] |
| $D_e'$ | $1.23 \times 10^{-4} \text{ cm}^2 \text{ day}^{-1}$ | [24, 25] |
| $D_T'$ | $8.64 \times 10^{-7} \text{ cm}^2 \text{ day}^{-1}$ | [25] |
| $\gamma_c'$ | $0.19 \times 10^{-10} \text{ mol.cell}^{-1}.\text{day}^{-1}$ | [19] |
| $\mu_c'$ | $0.0005 - 0.001 \text{ min}^{-1}$ | [31] |
| $k'$ | $2.3 \times 10^{-11} \text{ mol cell}^{-1} \text{ day}^{-1}$ | [32] |
| $\gamma_l'$ | $\frac{\gamma_c'}{3}$ | [33] |
| $\mu_l'$ | $0.24 - 2.4 \text{ day}^{-1}$ | [2] |
| $C^{*}'$ | $6 \text{ mmHg}$ | [21] |
| $C_H'$ | $5 \text{ mmHg}$ | [21] |
| $\lambda_{max}'$ | $4.95 \times 10^{-2} \text{ hour}^{-1}$ | [34] |
| $\mu_{max}'$ | $0.48 \text{ day}^{-1}$ | [28] |
| $L_N'$ | $20 \text{ mM}$ | [22] |
| $C_N'$ | $1 \text{ mmHg}$ | [23] |
| $\mu_d'$ | $0.024 \text{ day}^{-1}$ | [28] |
| $\lambda_{En}'$ | $0.05 - 0.33 \times 10^{-5} \text{ mM.cell}^{-1}.\text{day}^{-1}$ | Estimated |
| $\delta_E'$ | $0.16 - 0.5 \times 10^{-5} \text{ mM.cell}^{-1}.\text{day}^{-1}$ | Estimated |
| $L_{max}'$ | $12.8 - 14.5 \text{ mM}$ | Estimated |
| $\mu_E'$ | $0.8 - 0.98 \text{ day}^{-1}$ | Estimated |
| $\theta$ | 0 and 1 | — |

By dimensional analysis, the dimensionless values of all parameter for full model are given in Table 2.

**Table 2:** Dimensionless parameter values.

| Parameter | Value |
| --- | --- |
| $D_l$ | 0.25 |
| $D_e$ | $0.7 \times 10^{-4}$ |
| $D_T$ | $0.5 \times 10^{-6}$ |
| $\gamma_c$ | 1.59 |
| $\mu_c$ | $0.41 - 0.83$ |
| $k$ | 0.67 |
| $\gamma_l$ | 0.53 |
| $\mu_l$ | $0.14 - 1.44$ |
| $C^*$ | 0.6 |
| $\lambda_{max}$ | 0.77 |
| $\mu_{max}$ | 0.28 |
| $L_N$ | 0.5 |
| $C_H$ | 0.6 |
| $C_N$ | 0.1 |
| $\mu_d$ | 0.01 |
| $\theta$ | $0 - 1$ |
| $\lambda_{En}$ | $0.08 - 0.52$ |
| $\mu_E$ | $0.45 - 0.6$ |
| $L_{max}$ | $0.32 - 0.36$ |
| $\delta_E$ | $0.8 - 2.53$ |

## 4. Results

Unless we stated, numerical simulations for Eqs. (15)-(19) with boundary and initial conditions (Eqs. (20)-(24)) were generated using the parameter values in Table 2 for time-period 0 *< t < T*_*f*_ = 180. To simulate the model, we have used Forward time for temporal derivative and central difference for spatial derivative implicit scheme in the finite difference framework. The details description of discretizations for the governing equations are given in Appendix A. We also compute total tumor volume (*V*_*tot*_), total exosome (*E*_*tot*_) and lactate concentrations (*L*_*tot*_) as

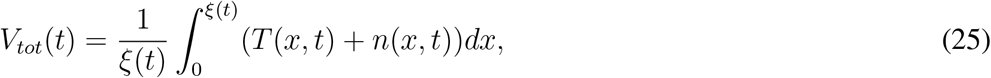

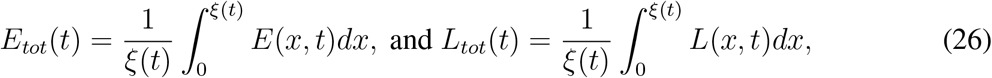

where *ξ*(*t*) is the length between the tumor center and the tumor tip point at each time step. *ξ*(*t*) is calculated using *T* (*ξ, t*) + *n*(*ξ, t*) = *ϵ* at each time point. Here, *ϵ* is a sufficiently small quantity, and in this study, we take *ϵ* = 10^*™*5^.

### 4.1. Effects of exosome feedback on tumor growth dynamics in oxygen-rich and oxygen-deprivation conditions

First, we explore exosome feedback on tumor growth under oxygen-rich (normoxic: *C > C*_*H*_) and oxygen-deprived (hypoxic: *C < C*_*H*_) conditions. We recall the parameter *θ* in Eq. (18), which controls the feedback of exosomes on tumor cell proliferation. Thus, *θ* = 0 represents the case in which this feedback is not accounted for, whereas *θ* = 1 denotes the case in which this feedback is included. Simulation results indicate that lactate is not secreted, and exosome levels for both *θ* = 0 and *θ* = 1 remain approximately the same over time under oxygen-rich conditions. It occurs because the positive feedback loop between lactate and exosome secretion does not occur in the absence of lactate under these culture conditions. From the time evolution of tumor cell density, we observe that the tumor density shows a negligible difference between the cases *θ* = 0 and *θ* = 1.

On the other hand, under oxygen-deprivation conditions, we observe that lactate levels are significantly higher for *θ* = 1 compared to *θ* = 0. As a result, exosome levels are also higher for *θ* = 1 than for *θ* = 0. However, the tumor cell density is significantly higher at *θ* = 1 than at *θ* = 0. Here, it is interesting to note that despite higher exosome levels under oxygen-deprived conditions for both cases, *θ* = 0 and *θ* = 1, the tumor volume is significantly lower than that observed under oxygen-rich culture conditions. This occurs because tumor cells undergo increased cell death in oxygen-deprived environments, driven by elevated lactate levels.

### 4.2. Tumor growth in symmetric cyclic hypoxia regimes

Recent experimental studies have shown that oxygen in the tumor microenvironment fluctuates between normoxic and hypoxic states over time, a phenomenon known as cyclic hypoxia. The time-scale of this fluctuation, i.e., the period of cyclic hypoxia, has a wide range in hours to days [23, 35]. This phenomenon is mimicked in the *in vitro* setting by tightly regulating oxygen levels in the culture medium. In most in vitro experiments done so far to recapitulate cyclic hypoxia, spatial heterogeneity has not been the focus, instead temporal fluctuations of oxygen levels have been imposed on cells [35, 36, 37, 38]. Hence, the mathematical form of the cyclic hypoxia protocol is given as follows

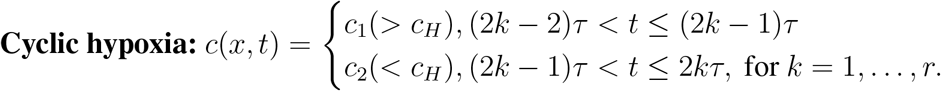

Where 2*rτ* = *T*_*final*_ is the total time of interest and 2*τ* is the period of cyclic hypoxia. The typical time frame of cyclic hypoxia is equal to one to five times the cell proliferation time scale, i.e., 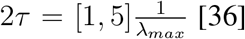; so, 2*τ* = [1.3, 6.5]. In this study, we choose the periods 2*τ* = 1.5, 2, 3, 4, 5 and 6. And such a choice ensures that the total durations of hypoxic and normoxic periods are equal at the end of the simulation.

Simulation results indicate that lactate levels increase significantly with an increasing period of cyclic hypoxia (Figure 3a). It occurs because the duration of hypoxic exposure increases as the period of cyclic hypoxia increases, leading to sustained lactate secretion. Additionally, exosome levels are observed to increase with longer periods of cyclic hypoxia, which is consistent with the elevated lactate levels that promote exosome production (Figure 3b). However, tumor volume decreases with increasing periods of cyclic hypoxia (Figure 3c). This reduction is attributed to the fact that high lactate levels induce a significant increase in cell death, which outweighs the exosome-driven tumor proliferation rate.

**Figure 3:**
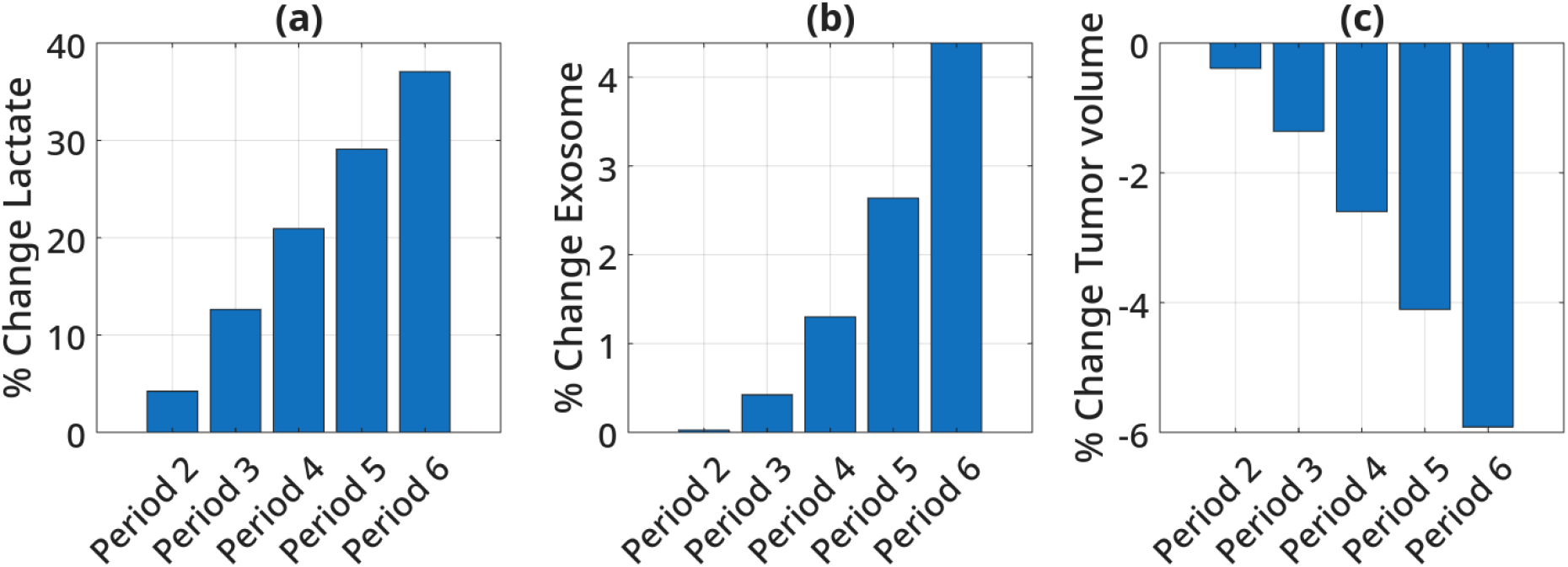
Time evolution of total (a) lactate, (b) exosome, and (c) tumor density in oxygen-rich and oxygen-deprived conditions. The solid line denotes the oxygen-deprivation case, and the dotted line denotes the oxygen-rich condition. The percentage change is calculated by comparing the total lactate, exosome concentrations, and tumor cell density in Period 1.5 with those in Periods 2, 3, 4, 5 and 6 at the end of the simulation.

### 4.3. Effects of hypoxic biasness on tumor growth

In vivo tumors, the durations of hypoxic and normoxic exposures during the periods of cyclic hypoxia are unequal. Therefore, the duration of normoxia and hypoxia is asymmetric. We define hypoxic bias as the percentage of hypoxic duration in a period of cyclic hypoxia. For example, hypoxic bias 0.75 corresponds to 75% of a period of cyclic hypoxia during hypoxic exposure. Hence, hypoxic bias 0.5 represents the symmetric duration of hypoxic and normoxic exposure in a period of cyclic hypoxia.

In this study, we consider the values of hypoxic bias to be 0.25, 0.4, 0.5, 0.6 and 0.75. To investigate the effects of hypoxic bias on lactate, exosomes, and tumor volume, we measure the percentage changes in lactate, exosomes, and tumor volume at the end of the simulation with respect to the values for the hypoxic bias of 0.5. Simulation results reveal that lactate levels drop for a small value of hypoxic bias (Figure 4a). Furthermore, as the hypoxic bias increases, lactate levels rise. It is because, at low hypoxic bias, the duration of hypoxic exposure is short; consequently, lactate production remains limited, while consumption occurs over a longer normoxic period. As the hypoxic bias increases, lactate accumulates to levels higher than those observed at a hypoxic bias of 0.5. Interestingly, increasing hypoxic bias also leads to a reduction in both exosome levels and overall tumor volume (Figures 4b and 4c). It may be attributed to enhanced tumor cell death induced by elevated lactate concentrations at higher hypoxic bias values.

**Figure 4:**
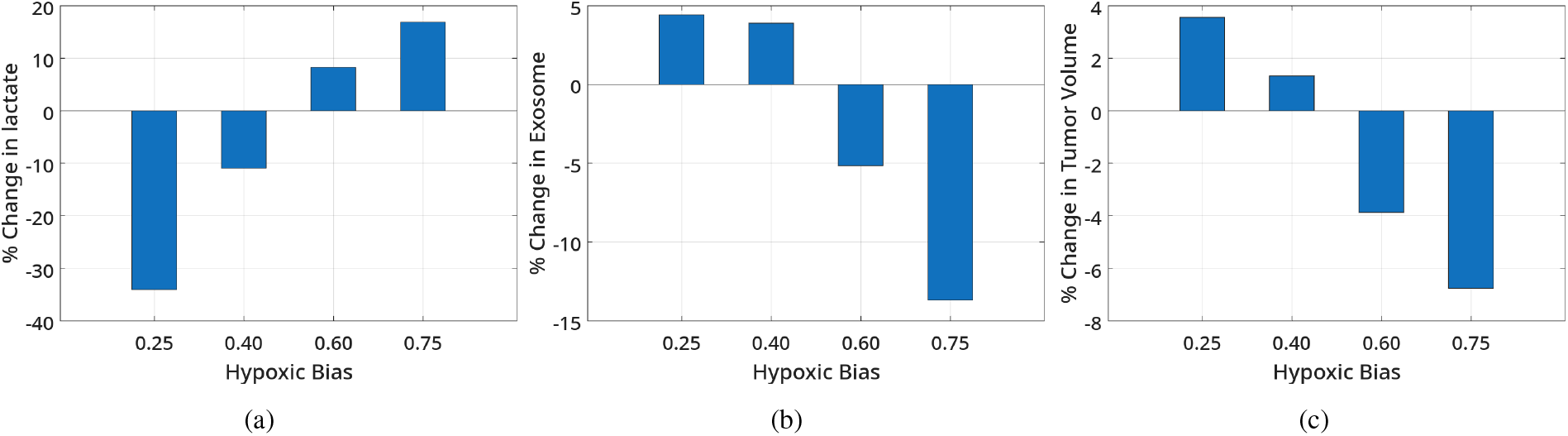
The effects of hypoxic bias on lactate, exosome and tumor volume for a period 5. The percentage changes of (a) lactate levels, (b) exosome levels and (c) tumor volume in hypoxic bias cases with respect to hypoxic bias 0.5 at the end of simulation.

In addition, similar trends are observed across different periods of cyclic hypoxia along the hypoxic bias axis. However, for hypoxic bias values below 0.5, lactate levels increase with increasing cycle periods. In contrast, for hypoxic bias values above 0.5, lactate levels decrease markedly as the period of cyclic hypoxia increases (Figure 5a). Exosome levels and tumor volume both decrease as the period value increases. When hypoxic bias exceeds 0.5, exosome levels and tumor volume are lower than those observed at a hypoxic bias of 0.5. In contrast, when hypoxic bias is below 0.5, these values are higher than at a bias of 0.5. At higher hypoxic bias, the percentage change in tumor volume during slow fluctuations also decreases (Figures 5b and 5c). These results suggest that tumor volume is reduced under conditions of increased hypoxic bias.

**Figure 5:**
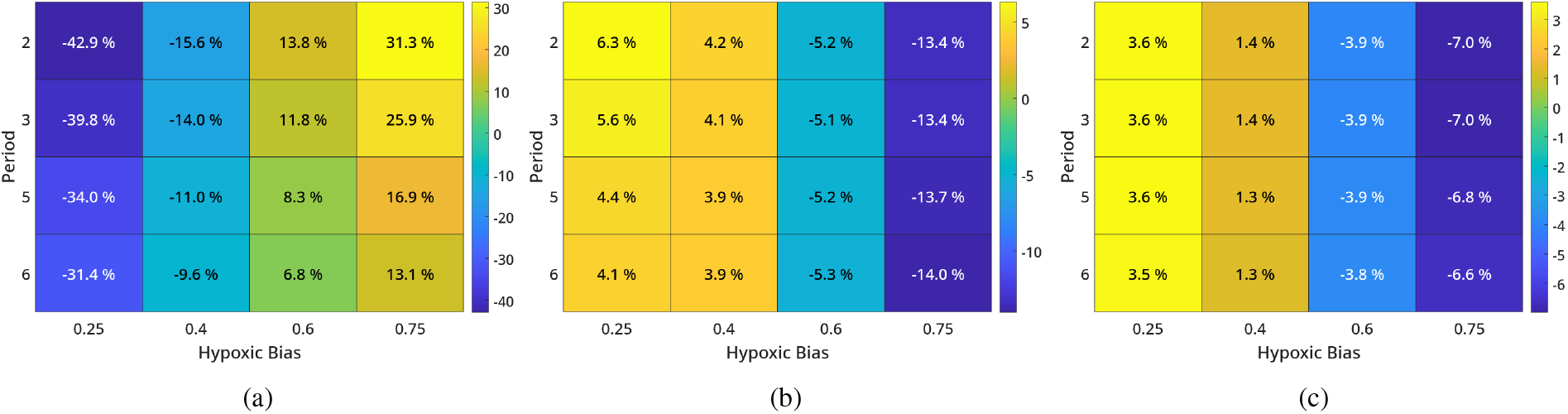
The effects of periods of cyclic hypoxia on lactate, exosome and tumor volume along with the different values of hypoxic bias. The heatmap is captured by formula 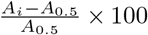 for *i* = 0.25, 0.4, 0.6 and 0.75 where *A* is lactate, exosome and tumor volume at the end of simulation.

### 4.4. Non-local interaction between lactate and exosome facilitates high tumor invasion under spatial heterogeneous oxygen environment

Spatial oxygen heterogeneity is a common feature of solid tumors. This heterogeneity arises from oxygen consumption by tumor cells and the limited diffusion of oxygen from blood vessels into the surrounding tissue [27, 35]. Simulation results display that spatial oxygen heterogeneity enables the coexistence of hypoxic and normoxic regions (Figure 6 a) as an oxygen gradient develops and falls below the hypoxic critical value. In hypoxic regions, lactate is produced, which provides positive feedback for exosome secretion. Tumor-derived exosomes accelerate tumor cell proliferation. Lactate produced in hypoxic regions diffuse into normoxic regions, where excessive accumulation may cause cell death. Hence, it is of interest to elucidate how lactate–exosome feedback influences tumor dynamics. Simulation results indicate that tumor cell density is higher when feedback from exosomes on tumor cell proliferation is accounted for (i.e., *θ* = 1) than when such feedback is not included (i.e., *θ* = 0), as expected (Figure 6 a).Moreover, it is observed that tumor cells in hypoxic regions benefit when *θ* = 1; however, this benefit decreases over time (Figure 6 c). It occurs because, for *θ* = 1, a larger hypoxic region develops over time compared to *θ* = 0 (Figure 6b). In the hypoxic region, lactate accumulation increases with time, and cell death eventually becomes dominant. In addition, we observe that tumor cells in the normoxic region benefit from exosome feedback on tumor cell proliferation after an initial transient period (Figure 6 b). It occurs because lactate-driven exosomes diffuse into the normoxic region and enhance tumor cell proliferation. In addition, we observe that the tumor size is larger for *θ* = 1 compared to *θ* = 0. The lactate–exosome axis not only affects the overall tumor size but also has a more pronounced impact on the size of the hypoxic core (Figure 6b). Hence, we conclude that the oxygen–lactate–exosome axis has the potential to increase intratumoral cellular heterogeneity.

**Figure 6:**
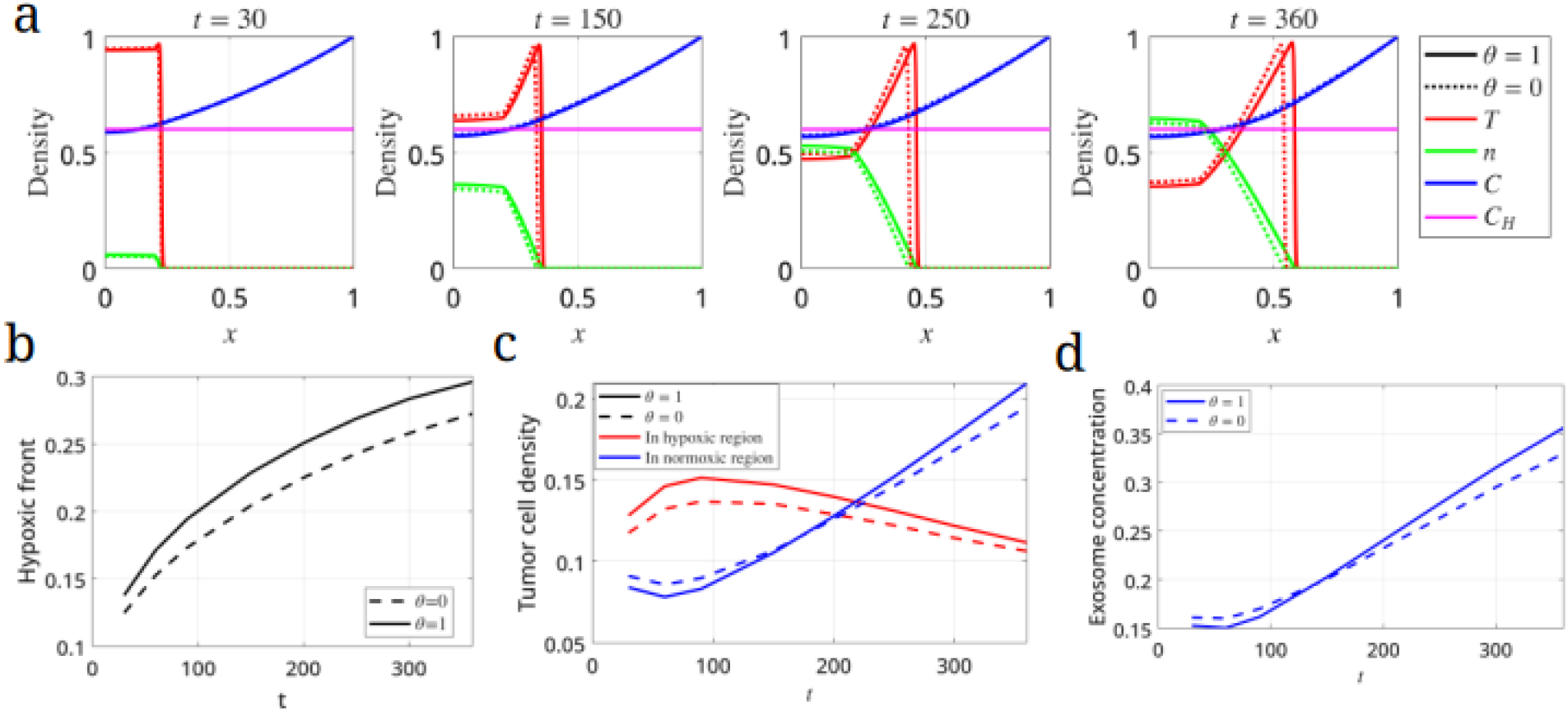
Spatial distribution of tumor cell, necrotic cell densities and oxygen concentration at *t* = 30, 150, 250 and 360 are displayed in (a) for both *θ* = 0 and 1. (b) The time evolution of hypoxic boundary is shown for both *θ* = 0 and 1. (c) Time evolution of tumor cell density in normoxic and hypoxic regions is displayed. (d) Exosome density change in normoxic region over time is portrayed. Here, the normoxic region is defined the region from 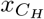 at where 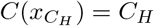 to tumor front *x*_*front*_.

## 5. Discussion

In this work, we have investigated tumor growth dynamics along the oxygen–lactate–exosome axis within a continuum mathematical modeling framework. Specifically, we have considered four distinct scenarios: (i) sustained oxygen-rich (normoxic) and oxygen-deprived (hypoxic) conditions; (ii) cyclic hypoxia characterized by periodic alternation between normoxic and hypoxic states with equal phase durations; (iii) cyclic hypoxia with unequal durations of the normoxic and hypoxic phases, i.e., environmental biasness; and (iv) spatial heterogeneity in oxygen distribution. The spatially homogeneous oxygen medium resembles the *in vitro* setup, while the heterogeneous oxygen environment represents the *in vivo* tumor. In addition, the time-course of oxygen levels in the growth medium mimics the *in vivo* event of cyclic hypoxia. Our proposed model is constructed based on partial differential equations (PDEs). The PDEs are derived based on the principle of mass conservation to describe the temporal dynamics of oxygen, lactate, exosomes, tumor cells, and necrotic cells. In this model, lactate is produced in hypoxic regions, where oxygen concentration falls below the hypoxic threshold, and is consumed in normoxic regions, where oxygen concentration exceeds the hypoxic threshold. Exosome secretion is modeled as a function of local lactate concentration, while tumor cell proliferation is regulated by local oxygen and exosome concentrations. Additionally, tumor cell death occurs when the local lactate concentration exceeds a critical value, and the oxygen concentration falls below a necrotic threshold.

Our numerical results demonstrated that lactate–exosome feedback on tumor growth is more significant in a chronic hypoxic environment than in a chronic normoxic environment. It arises because lactate is produced in a hypoxic environment, and this lactate enhances exosome production, thereby increasing tumor cell proliferation under chronic hypoxia. Experimental studies have shown that oxygen levels may fluctuate between normoxic and hypoxic states over a wide range of timescales, from minutes to days, a phenomenon known as cyclic hypoxia [39]. To examine the effects of cyclic hypoxia on the oxygen–lactate–exosome interplay and its role in tumor growth, we considered spatially homogeneous oxygen that varies temporally between normoxic and hypoxic levels. We observed that lactate and exosome levels increase as the period of cyclic hypoxia increases. However, it is interesting to note that tumor volume decreases as the period increases. This occurs because, over longer periods of cyclic hypoxia, the tumor produces ample amounts of lactate during hypoxic exposure, but it is not consumed extensively during the re-oxygenated phase; thus, lactate accumulates in the tumor microenvironment, leading to cell death. Moreover, the durations of hypoxic and normoxic exposures are not equal; rather, there is evidence that the environment is biased. Our results on hypoxic bias suggest that, with increasing hypoxic bias, lactate increases significantly, but due to the toxic effect of lactate, cells die; as a result, tumor volume and exosome levels decrease. *In vivo*, tumors exhibit spatially heterogeneous oxygen, and this heterogeneity arises mainly from oxygen consumption at the tumor periphery. In addition, our simulation results demonstrated that spatial oxygen heterogeneity enables tumor cells to benefit from lactate produced in hypoxic regions by increasing exosome production in normoxic regions. This ultimately increases tumor volume and invasiveness. Hence, it is concluded that the tumor is more aggressive when the oxygen–lactate–exosome feedback loop is activated in an oxygen-heterogeneous environment.

Our work suffers from several limitations. Here, we have only considered the tumor as having a mixed phenotype, with cells simultaneously proliferating and migrating. However, experimental studies have shown that tumors consist of heterogeneous phenotypic cells, where some specialize in proliferation and others in migration. This dichotomy is known as the go-or-grow hypothesis [40, 41, 42]. It would be interesting for future research to investigate how go-or-grow behavior is incorporated into the oxygen–lactate–exosome axis and to provide a more detailed picture of tumor progression. Since exosomes also play a key role in immune suppression [6], our modeling framework may be extended to investigate how tumor–immune interactions take shape in this complex feedback environment.

## Acknowledgment

GS gratefully acknowledges the Ministry of Education, Government of India, for research fellowship funding and the Indian Institute of Technology Guwahati for computational support.

## Data availability

This manuscript originates from Chapter 5 of the thesis entitled “A Mathematical Study on the Impacts of Oxygen Heterogeneity on Tumor Growth and EMT Dynamics” by Gopinath Sadhu. The thesis was submitted to the Indian Institute of Technology Guwahati on 3 April 2025 and defended on 17 June 2025. It was made available online on 20 July 2026.

The thesis can be accessed at the following link: https://gyan.iitg.ac.in/items/e256eb62-d231-4591-b7b7-1bad09a3261d

## Appendix A. Numerical method

The computational tissue domain [0, 1] was divided into (*N ™* 1) equal sub intervals 0 = *x*_0_ *< x*_1_ *< x*_2_ *<* … *< x*_*i™*1_ *< x*_*i*_ *< x*_*i*+1_ *<* … *< x*_*N™*2_ *< x*_*N™*1_ = 1 with uniform step length 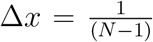. Δ*t* considers as the uniform time step length. Computation carries out up to final time *T*_*f*_ with the partition 0 *<* Δ*t <* 2Δ*t < · · · <* (*r ™* 1)Δ*t < r*Δ*t <* (*r* + 1)Δ*t < · · · < m*Δ*t*, where *m*+1 is the number of time steps. Note that 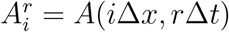, where *A* denotes *C, L, E, T* and *n*.

### Discretization of oxygen diffusion equation

Oxygen diffusion Eq. (15) is discretized using the central difference for spatial derivative and forward difference for temporal derivative. The discretized equation is obtained as,

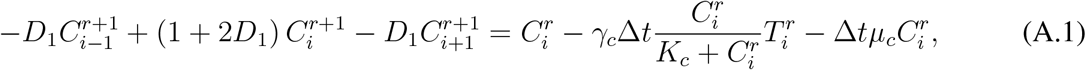

where *i* = 1, 2, 3, …, (*N ™* 3) and 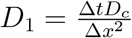.

At the points *i* = 0, (*N ™* 2), Eq. (A.1) became,

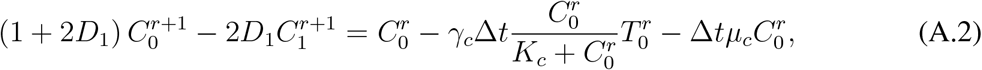

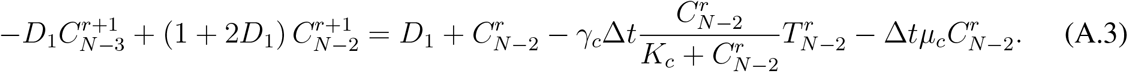

### Discretization of lactate diffusion equation

The lactate diffusion equation (16) is discretized as

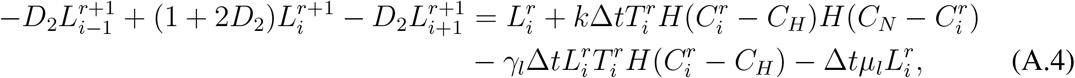

where 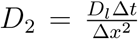. Since flux boundary conditions are available at the boundary, i.e., at *i* = 0, using ghost point criteria, Eq. (16) becomes

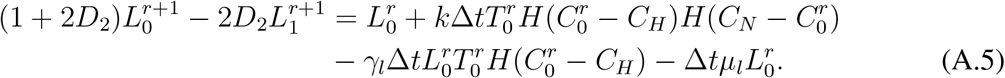

Also, at the boundary *i* = *N ™* 1, no flux condition is imposed. Using ghost point criteria, Eq. (16) become

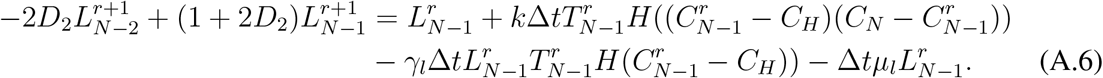

### Discretization of exosome diffusion equation

The exosome diffusion equation (17) is discretized as follow

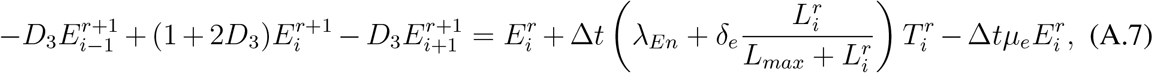

where 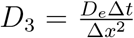 and *i* = 1, 2, …, *N ™* 2. At the boundary nodes i.e., at *i* = 0, *N ™* 1, Eq. (17) becomes

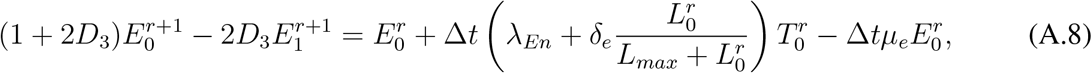

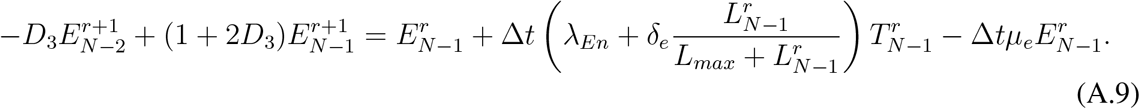

### Discretization of tumor cells density equation

The governing equation for the tumor cell density Eq. (18) is discretized as follows

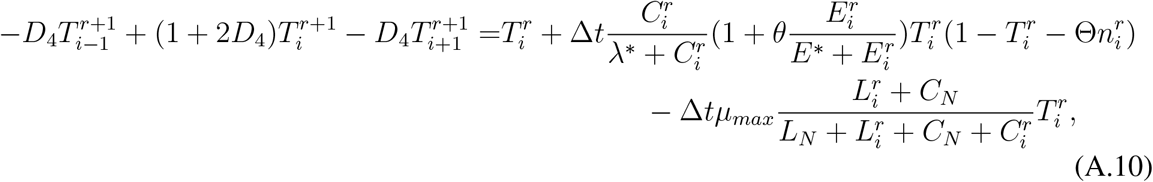

where 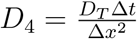 and *i* = 1, 2, …, *N ™* 2.

At the boundary nodes i.e., *i* = 0, *N ™* 1, the discretized form of Eq. 18 is given as

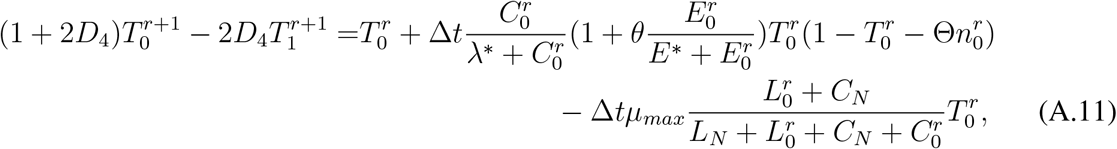

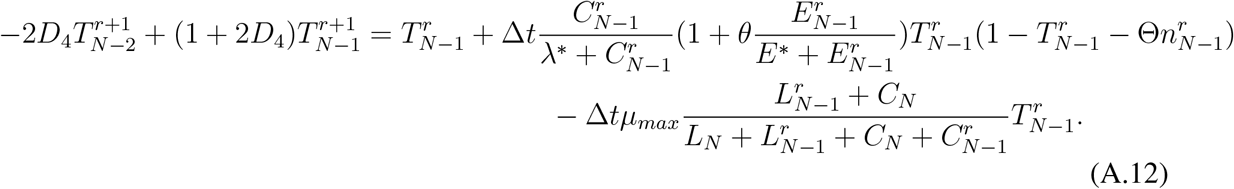

### Discretization of equation for necrotic cell dynamics

As the dynamics of the necrotic cell is governed by an ODE in Eq. (19) in non-dimensional form, we use the following formula

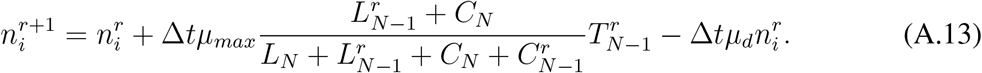

## Appendix B. Parameter estimation

When cells are cultured in vitro, nutrient, oxygen and lactate are maintained at constant, spatially uniform values and diffuse transport of exosomes can be neglected. In this case, Eq. (3) supplies

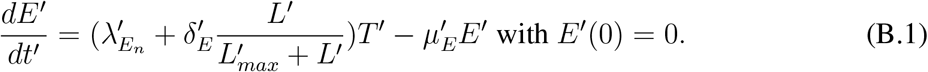

Its solution is given as

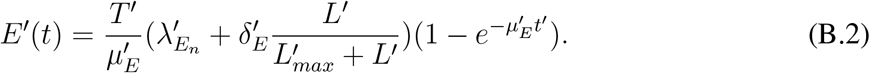

Here, the cells number changes in the experiment is neglected. Therefore, *T* ′ is constant. At 24 hours i.e., *t*′ = 1 day, Eq. (B.2) supplies

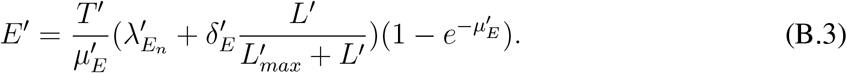

For notional conveniences, suppose 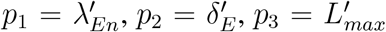 and 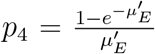. Eq. (B.3) supplies

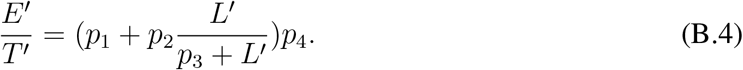

By fitting the data set (second row of Table B.3) from the melanoma cells to Eq. (B.4), we obtained the following parameter values: 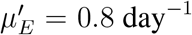, 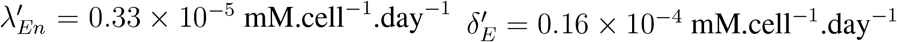 and 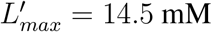.

**Table B.3:** Exosome level VS lactate concentration for B16-F10 melanoma and Hepa1-6 liver lines.

| L (mM) |  | 0 | 2 | 5 | 10 |
| --- | --- | --- | --- | --- | --- |
| Normalized $E$<br>(B16-F10) | | 0.07 | 0.13 | 0.14 | 0.22 |
| Normalized $E$<br>(Hepa1-6) | | 0.09 | 0.12 | 0.15 | 0.17 |

For the liver cells (the third row in Table B.3), the following parameter values: 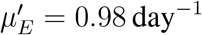, 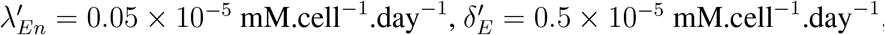, and 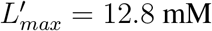.

## References

[1] D. Hanahan, R. A. Weinberg, The hallmarks of cancer, Cell 100 (1) (2000) 57–70.

[2] R. A. Gatenby, K. Smallbone, P. K. Maini, F. Rose, J. Averill, R. B. Nagle, L. Worrall, R. J. Gillies, Cellular adaptations to hypoxia and acidosis during somatic evolution of breast cancer, British journal of cancer 97 (5) (2007) 646–653.

[3] R. A. Gatenby, E. T. Gawlinski, A. F. Gmitro, B. Kaylor, R. J. Gillies, Acid-mediated tumor invasion: a multidisciplinary study, Cancer Research 66 (10) (2006) 5216–5223.

[4] M. Llibre, S. Kucuk, A. Gope, M. Certo, C. Mauro, Lactate: A key regulator of the immune response, Immunity 58 (3) (2025) 535–554.

[5] M. Logozzi, D. Mizzoni, D. F. Angelini, R. Di Raimo, M. Falchi, L. Battistini, S. Fais, Microenvironmental ph and exosome levels interplay in human cancer cell lines of different histotypes, Cancers 10 (10) (2018) 370.

[6] T. L. Whiteside, et al., Exosomes and tumor-mediated immune suppression, The Journal of Clinical Investigation 126 (4) (2016) 1216–1223.

[7] X. Yang, Y. Zhang, Y. Zhang, S. Zhang, L. Qiu, Z. Zhuang, M. Wei, X. Deng, Z. Wang, J. Han, The key role of exosomes on the pre-metastatic niche formation in tumors, Frontiers in Molecular Biosciences 8 (2021) 703640.

[8] E. Yang, X. Wang, Z. Gong, M. Yu, H. Wu, D. Zhang, Exosome-mediated metabolic reprogramming: the emerging role in tumor microenvironment remodeling and its influence on cancer progression, Signal transduction and targeted therapy 5 (1) (2020) 242.

[9] J.-L. Qu, X.-J. Qu, M.-F. Zhao, Y.-E. Teng, Y. Zhang, K.-Z. Hou, Y.-H. Jiang, X.-H. Yang, Y.-P. Liu, Gastric cancer exosomes promote tumour cell proliferation through pi3k/akt and mapk/erk activation, Digestive and liver disease 41 (12) (2009) 875–880.

[10] J. A. Bull, H. M. Byrne, The hallmarks of mathematical oncology, Proceedings of the IEEE 110 (5) (2022) 523–540.

[11] H. M. Byrne, J. R. King, D. S. McElwain, L. Preziosi, A two-phase model of solid tumour growth, Applied Mathematics Letters 16 (4) (2003) 567–573.

[12] G. Sadhu, A mathematical investigation of exosome and lactate levels interplay in an in vitro and in vivo tumors, Bulletin of Mathematical Biology 88 (1) (2026) 4.

[13] M. Al-Husari, S. D. Webb, Regulation of tumour intracellular ph: A mathematical model examining the interplay between h+ and lactate, Journal of Theoretical Biology 322 (2013) 58–71.

[14] A. S. Moffett, Y. Deng, H. Levine, Modeling the role of immune cell conversion in the tumor-immune microenvironment, Bulletin of Mathematical Biology 85 (10) (2023) 93.

[15] D. I. Wallace, X. Guo, Properties of tumor spheroid growth exhibited by simple mathematical models, Frontiers in oncology 3 (2013) 51.

[16] C. Colson, P. K. Maini, H. M. Byrne, Investigating the influence of growth arrest mechanisms on tumour responses to radiotherapy, Bulletin of Mathematical Biology 85 (8) (2023) 74.

[17] C. Colson, H. M. Byrne, P. K. Maini, Combining mechanisms of growth arrest in solid tumours: a mathematical investigation, Bulletin of Mathematical Biology 84 (8) (2022) 80.

[18] H. Greenspan, Models for the growth of a solid tumor by diffusion, Studies in Applied Mathematics 51 (4) (1972) 317–340.

[19] M. Shamsi, M. Saghafian, M. Dejam, A. Sanati-Nezhad, Mathematical modeling of the function of warburg effect in tumor microenvironment, Scientific Reports 8 (1) (2018) 8903.

[20] J. J. Casciari, S. V. Sotirchos, R. M. Sutherland, Mathematical modelling of microenvironment and growth in emt6/ro multicellular tumour spheroids, Cell Proliferation 25 (1) (1992) 1–22.

[21] P. Sonveaux, F. Végran, T. Schroeder, M. C. Wergin, J. Verrax, Z. N. Rabbani, C. J. De Saedeleer, K. M. Kennedy, C. Diepart, B. F. Jordan, et al., Targeting lactate-fueled respiration selectively kills hypoxic tumor cells in mice, The Journal of Clinical Investigation 118 (12) (2008) 3930–3942.

[22] G. Sadhu, D. C. Dalal, A mathematical study of the interaction between oxygen and lactate in an in vivo and in vitro tumor, International Journal of Biomathematics (2024) 2450138.

[23] G. Sadhu, D. C. Dalal, Effects of non-linear interaction between oxygen and lactate on solid tumor growth under cyclic hypoxia, Bulletin of Mathematical Biology 87 (3) 41.

[24] X. Lai, A. Friedman, Exosomal mirs in lung cancer: a mathematical model, PLoS One 11 (12) (2016) e0167706.

[25] X. Lai, A. Friedman, Exosomal microrna concentrations in colorectal cancer: A mathematical model, Journal of Theoretical Biology 415 (2017) 70–83.

[26] C. Michiels, C. Tellier, O. Feron, Cycling hypoxia: A key feature of the tumor microenvironment, Biochimica et Biophysica Acta (BBA)-Reviews on Cancer 1866 (1) (2016) 76–86.

[27] G. Sadhu, K. Yadav, S. S. Ghosh, D. Dalal, The impact of oxygen distribution on the tumor necrotic region: A two-phase model, Acta Biotheoretica 74 (2) (2026) 9.

[28] M. Chaplain, G. Lolas, Mathematical modelling of cancer invasion of tissue: dynamic heterogeneity., Networks and Heterogeneous Media 1 (3) (2006) 399–439.

[29] D. R. Grimes, C. Kelly, K. Bloch, M. Partridge, A method for estimating the oxygen consumption rate in multicellular tumour spheroids, Journal of The Royal Society Interface 11 (92) (2014) 20131124.

[30] M. Luo, J. Zhu, J. Ren, Y. Tong, L. Wang, S. Ma, J. Wang, Lactate increases tumor malignancy by promoting tumor small extracellular vesicles production via the gpr81-camp-pka-hif-1α axis, Frontiers in Oncology 12 (2022) 1036543.

[31] C. K. Griffith, C. Miller, R. C. Sainson, J. W. Calvert, N. L. Jeon, C. C. Hughes, S. C. George, Diffusion limits of an in vitro thick prevascularized tissue, Tissue Engineering 11 (1-2) (2005) 257–266.

[32] M. Papadogiorgaki, M. G. Kounelakis, P. Koliou, M. E. Zervakis, A glycolysis-based in silico model for the solid tumor growth, IEEE journal of biomedical and health informatics 19 (3) (2014) 1106–1117.

[33] J. B. McGillen, C. J. Kelly, A. Martínez-González, N. K. Martin, E. A. Gaffney, P. K. Maini, V. M. Pérez-García, Glucose–lactate metabolic cooperation in cancer: Insights from a spatial mathematical model and implications for targeted therapy, Journal of Theoretical Biology 361 (2014) 190–203.

[34] R. Venkatasubramanian, M. A. Henson, N. S. Forbes, Incorporating energy metabolism into a growth model of multicellular tumor spheroids, Journal of Theoretical Biology 242 (2) (2006) 440–453.

[35] G. Sadhu, H. M. Byrne, D. C. Dalal, The impact of oxygen heterogeneity on epithelial-to-mesenchymal transitions: a numerical study, Journal of Mathematical Biology 92 (1) (2026) 15.

[36] G. Sadhu, P. Jain, J. T. George, M. K. Jolly, A phenotype-structured pde framework for investigating the role of hypoxic memory on tumor invasion under cyclic hypoxia, Bulletin of Mathematical Biology (2026) 2025–10.

[37] G. L. Celora, S. B. Bader, E. M. Hammond, P. K. Maini, J. M. Pitt-Francis, H. M. Byrne, A dna-structured mathematical model of cell-cycle progression in cyclic hypoxia, Journal of Theoretical Biology 545 (2022) 111104.

[38] G. L. Celora, R. Nixson, J. M. Pitt-Francis, P. K. Maini, H. M. Byrne, Characterising cancer cell responses to cyclic hypoxia using mathematical modelling, Bulletin of Mathematical Biology 86 (12) (2024) 145.

[39] K. Saxena, M. K. Jolly, Acute vs. chronic vs. cyclic hypoxia: their differential dynamics, molecular mechanisms, and effects on tumor progression, Biomolecules 9 (8) (2019) 339.

[40] H. Hatzikirou, D. Basanta, M. Simon, K. Schaller, A. Deutsch, ‘go or grow’: the key to the emergence of invasion in tumour progression?, Mathematical Medicine and Biology: A Journal of the IMA 29 (1) (2010) 49–65.

[41] R. Thiessen, M. Conte, T. L. Stepien, T. Hillen, Go-or-grow models in biology: a monster on a leash, Journal of Mathematical Biology 91 (5) (2025) 58.

[42] C. Falcó, R. M. Crossley, R. E. Baker, Travelling waves in a minimal go-or-grow model of cell invasion, Applied Mathematics Letters 158 (2024) 109209.

